# Evidence for a role of human blood-borne factors in mediating age-associated changes in molecular circadian rhythms

**DOI:** 10.1101/2023.04.19.537477

**Authors:** Jessica E. Schwarz, Antonijo Mrčela, Nicholas F. Lahens, Yongjun Li, Cynthia T. Hsu, Gregory Grant, Carsten Skarke, Shirley L. Zhang, Amita Sehgal

**Author notes:** Email: Amita Sehgal,; Shirley L. Zhang. Author Contributions: Conceptualization, J.E.S. and A.S.; Methodology, J.E.S, A.M., N.F.L., Y.L., C.T.H., G.G., C.S., S.L.Z, and A.S.; Formal Analysis, J.E.S, S.L.Z, A.M., N.F.L., Y.L., and C.T.H; Investigation, J.E.S, S.L.Z., and C.S.; Writing – Original Draft Preparation, J.E.S. and A.S.; Writing – Review & Editing, J.E.S, A.M., N.F.L., Y.L., C.T.H., G.G., C.S., S.L.Z, and A.S.; Visualization, J.E.S., A.M, N.F.L.; Supervision, S.L.Z and A.S.

## Abstract

Aging is associated with a number of physiologic changes including perturbed circadian rhythms; however, mechanisms by which rhythms are altered remain unknown. To test the idea that circulating factors mediate age-dependent changes in peripheral rhythms, we compared the ability of human serum from young and old individuals to synchronize circadian rhythms in culture. We collected blood from apparently healthy young (age 25-30) and old (age 70-76) individuals at 14:00 and used the serum to synchronize cultured fibroblasts. We found that young and old sera are equally competent at initiating robust ∼24h oscillations of a luciferase reporter driven by clock gene promoter. However, cyclic gene expression is affected, such that young and old sera promote cycling of different sets of genes. Genes that lose rhythmicity with old serum entrainment are associated with oxidative phosphorylation and Alzheimer’s Disease as identified by STRING and IPA analyses. Conversely, the expression of cycling genes associated with cholesterol biosynthesis increased in the cells entrained with old serum. Genes involved in the cell cycle and transcription/translation remain rhythmic in both conditions. We did not observe a global difference in the distribution of phase between groups, but found that peak expression of several clock-controlled genes (*PER3, NR1D1, NR1D2, CRY1, CRY2,* and *TEF*) lagged in the cells synchronized *ex vivo* with old serum. Taken together, these findings demonstrate that age-dependent blood-borne factors affect circadian rhythms in peripheral cells and have the potential to impact health and disease via maintaining or disrupting rhythms respectively.

## INTRODUCTION

Circadian rhythms are known to regulate homeostatic physiology including sleep:wake, hormone production and body temperature, and their dysregulation with aging is accompanied by adverse health consequences^1–3^, raising the possibility that health decline with age is caused in part by circadian dysfunction. Although the mechanisms responsible for age effects on circadian rhythms are unknown, signals from the central clock in the suprachiasmatic nucleus (SCN) dampen with age^3–5^ and rhythms change in peripheral tissues in different ways^5,6^. Here we aimed to develop a cell culture model to study the effect of aging on human rhythms of peripheral tissues. Given that serum can reset the clock in peripheral fibroblasts^7^, we questioned the extent to which serum factors normally contribute to peripheral rhythms of gene expression and how they might affect rhythms with age, given that blood-borne factors can influence other aspects of aging^8^.

Using an established clinical study paradigm that sampled blood at 14:00^9^, we collected blood from young (age 25-30) and old (age 70-76) apparently healthy individuals with behavioral and physiological outputs quantified by wearable devices, and we tested the hypothesis that age-dependent factors in the sera affect circadian rhythms in cultured fibroblasts. In support of this hypothesis, we show here that genes associated with oxidative phosphorylation and mitochondrial functions lose rhythmicity in fibroblasts exposed to aged serum factors. We also find evidence of reduced entrainment in terms of altered expression of several molecular clock genes (*PER3, NR1D1, NR1D2, CRY1, CRY2* and *TEF*) when synchronized with aged serum. These findings suggest that age-related changes in blood borne factors contribute to impaired circadian physiology and the associated disease risks.

## RESULTS

We enrolled 8 old and 7 young human subjects (Fig.1A, Tables S1 and Fig. S2), whose demographics are listed in Table S1. The behavioral and physiological assessments confirmed entrainment to the light-dark conditions local to the East Coast of the US. This is, for example, evident in the diurnal rhythms observed for physical activity (Fig. 1C top, vector magnitude two-way ANOVA for time-of-day *q*=3.3E-06), light exposure (lux two-way ANOVA for time-of-day *q*=0.01), heart rate (bpm two-way ANOVA for time-of-day *q*=0.016) and sympathetic and parasympathetic nervous system indices derived from the Kubios heart rate variability analysis (two-way ANOVA for time-of-day *q*=0.099 and *q*=0.063, respectively).

**Fig 1.**
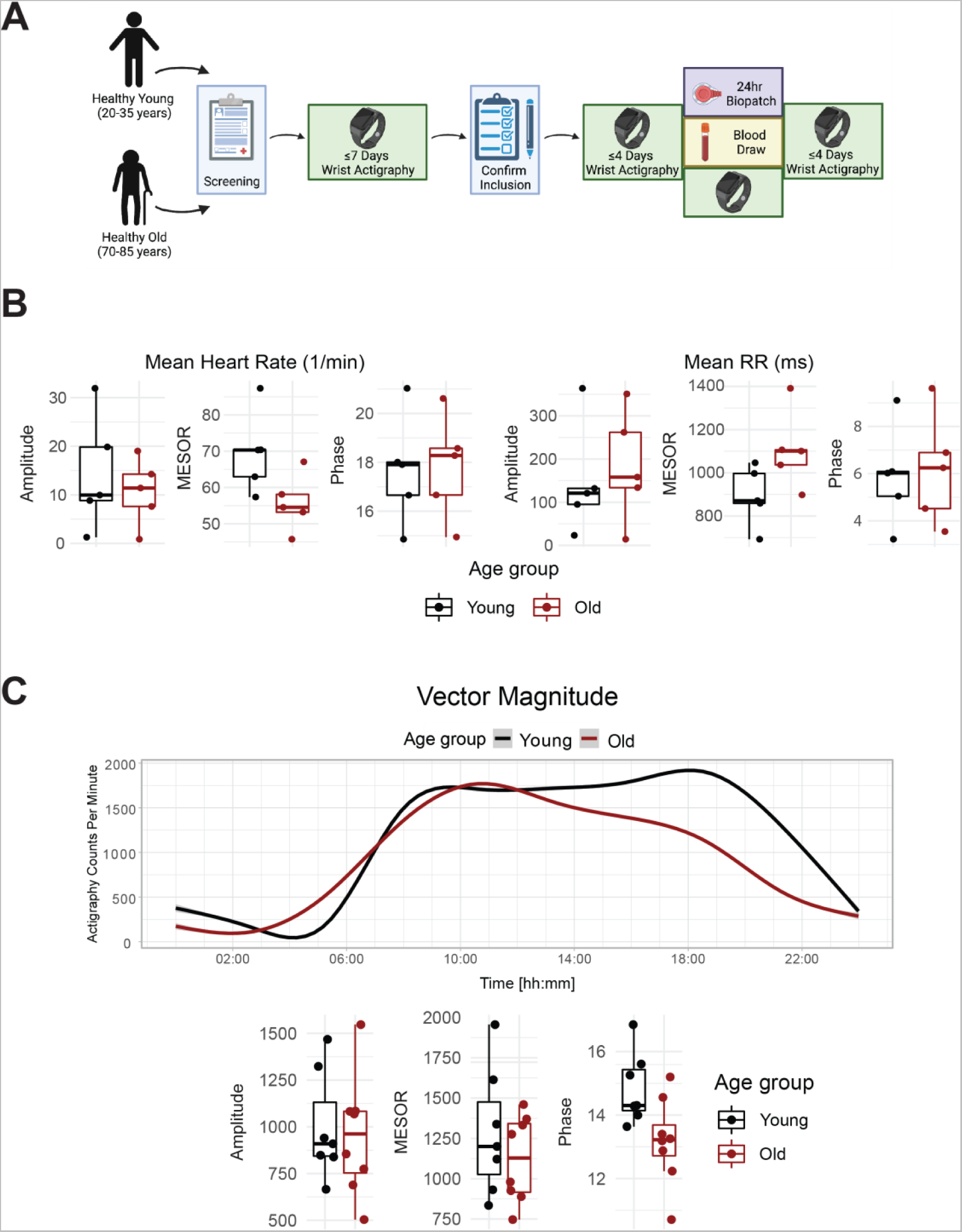
Healthy elderly individuals tend to have lower heart rate, increased heart rate variability, and phase advanced activity patterns relative to young subjects. Experimental protocol for enrolment and monitoring of human subjects (A, made with BioRender.com). As assessed by Zyphyr BioPatch, electrocardiogram (EKG) measurements suggest that MESOR of average heart rate decreases with age (p=0.056) and that MESOR of heart rate variability increases with age (p=0.056). N=5 per group. MESOR and amplitude were tested using two-sided Wilcoxon rank sum exact test, while phase was tested by Kuiper’s two-sample test (B). Average activity counts across three axes (vector magnitude), as recorded by the actigraph device plotted throughout the day (top) and analyzed for circadian rhythm (bottom) (C). While the amplitudes of activity did not differ between the age groups, the older individuals trended towards an early phase (p∼0.055, Kupier’s two-sample test) compared to young individuals. N=7 for young and N=8 for old. Lines in the top panel of C are smoothed means (fit with penalized cubic regression splines) for data from each age group. Dots in the bottom panel of C are subject-level cosinor parameter estimates derived from cosinor fits to the actigaraphy data. Boxplot midlines correspond to median values, while the lower and upper hinges correspond to the first and their quartiles, respectively. Boxplot whiskers extend to the smallest/largest points within 1.5 * IQR (Inter Quartile Range) of the lower/upper hinge.

Age-specific differences emerged for several outputs. The standard deviation of instantaneous heart rate values was significantly lower in the old compared to young (two-way ANOVA for age group q=0.042). As expected, the peak clock time (acrophase) of physical activity (triaxial accelerometry integrated as vector magnitude) was phase-advanced among old compared to young subjects, as was the midsleep time calculated from the MCTQ (Fig S1B), though not at a statistically significant level (p=0.055). On average, a lower rhythm-adjusted mean (mesor) heart rate of 55.8±3.5 bpm was found in old compared to 69.7±5.0 bmp in young, along with a higher heart rate variability (RR intervals) of 1106.7±80.5 ms compared to 893.2±61.8 ms, respectively (Fig 1B). Old subjects displayed lower activity in the sympathetic nervous system (SNS), and higher activity in the parasympathetic nervous system (PNS) compared to young (Fig. S1A), with no significant difference in cortisol levels (Fig. S1C). The overall high degree of variability rendered the cardiovascular trends, however, statistically not significant.

As noted above, blood was collected from these old/young individuals and the serum was used to synchronize BJ-5TA fibroblasts stably transfected with a *BMAL1-luciferase* construct^7,9–11^. Circadian rhythms were assessed by luciferase assay^10^ over 4 days (Fig 2A). We conducted this study in a fibroblast line in order to build upon age-related changes found in a previous study which used human fibroblast lines^9^. An additional rationale for using this line is that it derives from normal human tissue, providing an advantage in studying normal physiology when compared to the commonly used U2OS line which is a genomically unstable osteosarcoma with chromosomal abnormalities^12^. No significant differences were observed in the period, amplitude, and phase of the *BMAL1-luciferase* rhythm with young versus old serum treatment (Fig S3).

**Fig 2.**
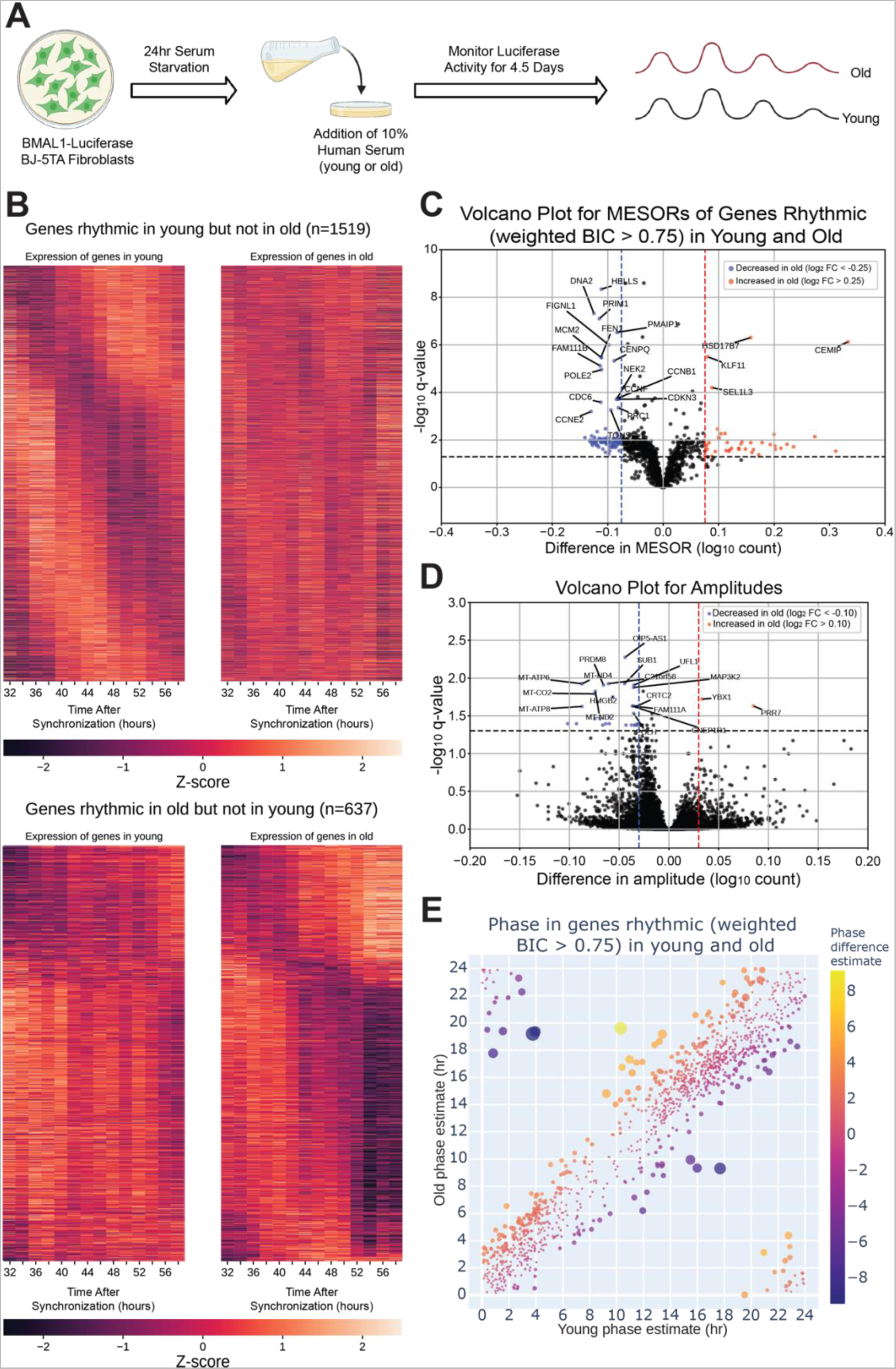
The circadian transcriptome is differentially affected by entrainment with sera from young and old human subjects. A visual representation of the serum starvation-serum addition protocol to synchronize the *BMAL1-luciferase* BJ-5TA fibroblasts **(A**, made with BioRender.com). When comparing the circadian transcriptome entrained by either young or old sera (n=4 sera per group), 1519 genes lost rhythmicity with age (B, top) while only 637 genes gained rhythmicity with age (B, bottom). Weighted BIC criterion with a threshold of 0.75 was used to assess rhythmicity **(B)**. The number of genes rhythmic in young and old (according to weighted BIC > 0.75 criterion) that show a detectable change in MESOR (q < 0.05 criterion for MESOR difference) is 568, with MESOR increasing in 163 and decreasing in 405 genes. Out of these, 40 genes with increased MESOR (red) and 98 genes with decreased MESOR (blue) also satisfy the condition |log_2_FC| > 0.25 **(C)**. We were only able to detect change in amplitude for a small number of genes, 39 genes had decreased amplitude in old and 2 had increased amplitude using CircaCompare. Only 30 genes with decreased amplitude also satisfy the condition |log_2_ FC| > 0.1 (blue) **(D)**. For phase, using a test provided by CircaCompare, under q < 0.05 cutoff for age-related phase differences in genes rhythmic in young and old (with BIC>0.75 cycling criteria), we detected 20 genes with advanced phase, and 34 with delayed phase **(E)**.

To assess serum effects on circadian gene expression, we first performed RNA-seq on fibroblasts synchronized with serum from a single old or single young individual at two hour intervals and found that, compared to the first day of synchronization, the second day (36-58 hours after synchronization) showed greater differences in MESORs between young and old serum-treated groups (Fig. S4). This is not surprising because the first day includes acute responses of the fibroblasts to serum, which can mask circadian rhythms, and so the second day is expected to reveal differences in endogenous rhythms between samples. Day 2 also revealed different phases of cyclic expression between young and old subjects for a larger number of genes. We proceeded to collect fibroblasts synchronized by sera from eight different subjects (four old, four young with two male and two female per group) at two-hour intervals, from 32 to 58 hours post serum addition. This added an extra two timepoints to the second day to facilitate the calculation of rhythmicity. Using CircaCompare^13^ and a weighted Bayesian Information Criterion (BIC)>0.75 cutoff, a significant number of genes were found to lose (1519 genes) or gain (637 genes) rhythmicity with age, underscoring the impact of age on the ability of serum to synchronize circadian rhythms, while 1209 genes were rhythmic in both groups (Fig 2B, 3A). Additionally, we used CircaCompare to estimate MESORs, amplitudes, and phases of gene oscillatory patterns in young and old groups, and to compare these cosinor parameters between groups (Fig 2C-D). Of the genes that were rhythmic in both groups, many also showed changes with age. For instance, 568 cyclically expressed genes showed a change in MESOR with age (q < 0.05 for MESOR differences), with MESOR values increasing for 163 genes and decreasing for 405 genes in the old serum-treated samples (Fig 2C, S5, right). Using q-values provided by CircaCompare we were able to detect changes in amplitude (Fig 2D, S5, left) for only a small number of genes (39 genes had decreased amplitude in old and 2 had increased amplitude). Using CircaCompare and a q < 0.05 cutoff for phase differences in genes rhythmic in young and old, we detected 20 genes with advanced phase, and 34 with delayed phases (Fig 2E). However, it is important to note that in these analyses low p-values can be driven by large sample-size such as in Fig S5 where distributions of MESORs and amplitudes across genes are assessed such that each gene represents a sample.

As mentioned, we found several examples of transcripts that cycle with young serum synchronization and lose rhythmicity (ex. TCF4), decrease in amplitude (ex. HMGB2), phase shift (ex. TEF), or change MESOR (ex. HSD17B7) with aged serum (Fig. S6). Search Tool for the Retrieval of Interacting Genes (STRING)^14^ and Ingenuity Pathway Analysis (IPA)^15^ were used for functional genomics. Both approaches indicate a maintenance of cycling of cell cycle genes with young and old serum synchronization, and a loss of rhythmicity of genes associated with oxidative phosphorylation in the aged serum (Fig 3A and S7). STRING analysis revealed that the dominant pathways associated with genes rhythmic in both young and old conditions, cell cycle and DNA replication, demonstrate a decrease in MESOR with old serum. For instance, checkpoint control and chromosomal replication pathway associated genes were expressed cyclically in both young and old conditions; however, several chromosomal replication pathway genes exhibit decreased MESORs in the aged sera (Fig S7 D,E, Table S3). MESORs of steroid biosynthesis genes, particularly those relating to cholesterol biosynthesis, were increased in the old sera condition (Fig 3A, Table S4).

**Fig 3.**
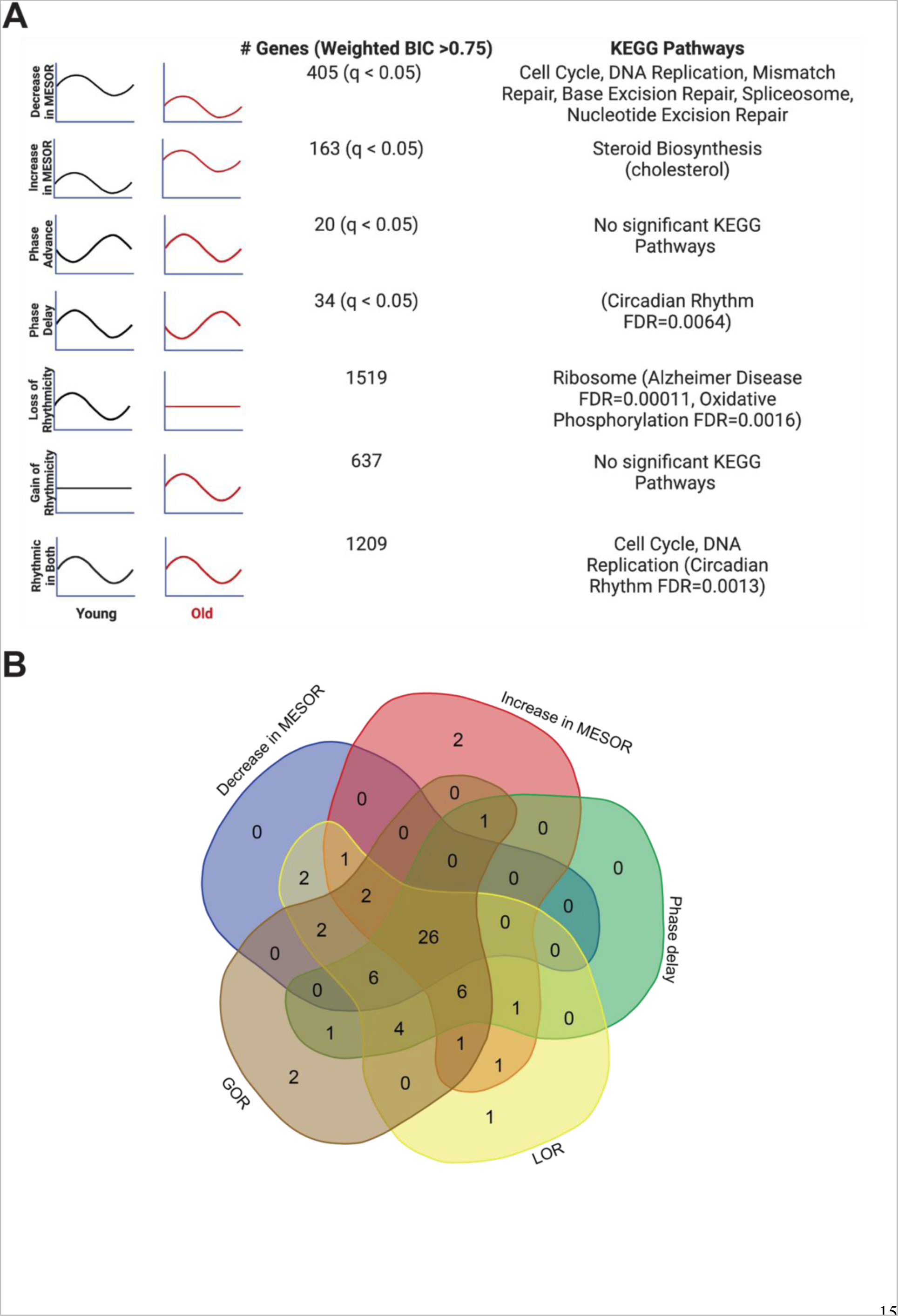
Each type of circadian change is associated with different KEGG pathways by STRING analysis, but a similar set of transcription factors identified by LISA. Entrainment of the fibroblasts in culture with old serum significantly altered the circadian transcriptome despite use of the same cells in both young and old conditions, suggesting the aged serum itself affects the regulation of specific pathways in the cell. Significant KEGG Pathways (FDR <1x10^-4^ unless specified) are indicated next to each category of genes. Age related pathways such as Alzheimer’s Disease/oxidative phosphorylation are associated with a loss of rhythms in the old condition. Cell cycle and DNA replication pathways remain rhythmic in the old serum condition, but cycle with a decreased MESOR. Rhythmic genes were determined using CircaCompare. Rhythmicity was determined using the weighted BIC > 0.75 criterion while p-values for difference in MESOR and phase were determined using CircaCompare **(A)**. LinC similarity analysis (LISA), based on known transcription factor binding in fibroblasts and RNA expression, suggests that 59 total transcription factors (q<0.05) show significant changes in activity in conjunction with the following cycling phenotypes: decreased in MESOR, increased in MESOR, phase delay, gain of rhythmicity, and loss of rhythmicity as were defined above **(B)**.

Loss of rhythms in oxidative phosphorylation suggests mitochondrial dysfunction with age. By STRING analysis, 24 out of the 26 genes associated with oxidative phosphorylation overlap with the Alzheimer’s Disease KEGG pathway highlighting the disease relevance of this class of genes that loses cycling in older individuals. Given that Alzheimer’s pathology is closely associated with the accumulation of oxidative stress^16^, it is possible that loss of cycling contributes to oxidative damage. An additional 31 genes in the Alzheimer’s Disease KEGG pathway lose rhythmicity with aged serum; these include amyloid precursor protein (APP) and apolipoprotein E (APOE). APP is by definition the precursor of amyloid beta, which accumulates in AD^17^. While APOE plays an important role in lipid transport, specific variants of this gene are strongly associated with AD risk as APOE interacts with amyloid beta in amyloid plaques, a hallmark of the disease ^18^.

To determine whether aged serum modifies the activity of specific transcription factors, we performed an epigenetic Landscape In Silico deletion analysis (LISA), a computational tool designed to predict changes in transcription factor activity based on gene expression. Using a comprehensive database of known transcription factor binding profiles of fibroblasts, we identified altered gene expression corresponding to 59 total transcription factors (q<0.05) in our young-versus-old cycling dataset. Potential changes in the activity of these transcription factors are associated with the following five categories of cycling phenotypes: decreased MESOR, increased MESOR, phase delay, gain of rhythmicity, and loss of rhythmicity, as were defined above (Fig 3B). 26 transcription factors were represented in all groups, and included those implicated in diseases such as cancer (MYC, TP53, YAP1, RBL2, E2F7, BRD4, MXl1), inflammation (CEBPB, BHLHE40), oxidative stress (NFE2L2), and neurological conditions (NEUROG2, SUMO2).

Lastly, several clock genes showed differences in expression with the aged serum, most notably genes in the Circadian Rhythm KEGG pathway (Fig 4). In particular, expression of *CRY1*, *CRY2*, *NR1D1*, *NR1D2*, *PER3*, and *TEF* was significantly phase delayed after synchronization with old serum compared to young (Fig 4). Importantly, the RNA-seq did not reveal a difference in the phase or amplitude of *BMAL1* expression with age, supporting the validity of our *BMAL1-luciferase* findings, although the MESOR significantly increased with age.

**Fig 4.**
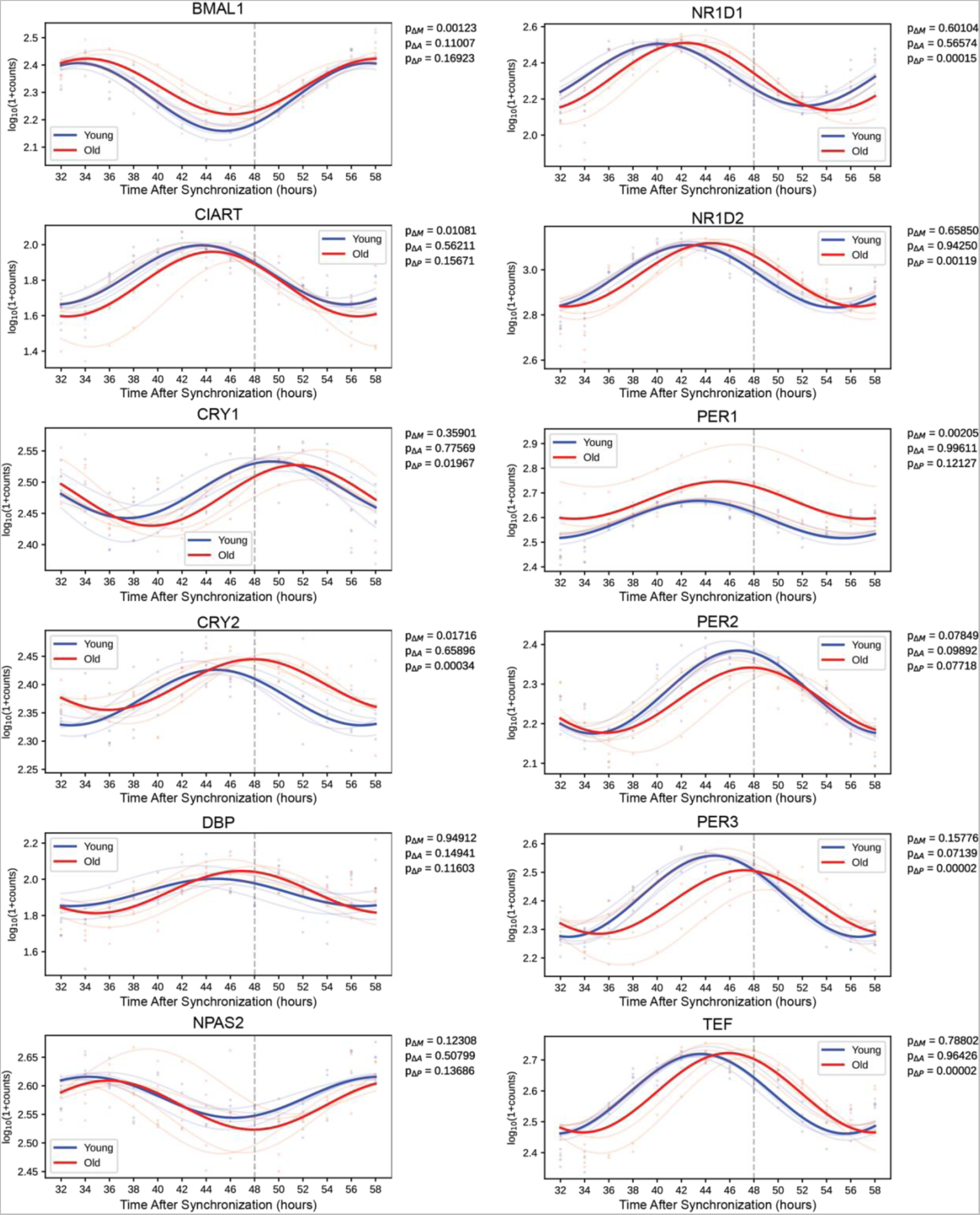
Synchronizing with old serum phase delays the expression profile of several core clock genes. Traces of molecular clock mRNA transcripts, the average curve (bold) of results of individual sera (faded). P-values for the difference in MESOR (P_ΔM), amplitude (P_ΔA), and phase (P_ΔP) are shown for the comparison of young and old conditions. Several clock genes are significantly phase delayed (*CRY1, CRY2, NR1D1, NR1D2, PER3, TEF*) in response to synchronization with old serum. While *BMAL1* is not phase delayed, the MESOR significantly increases with age. N=4 subjects per timepoint for both young and old groups.

## DISCUSSION

Studies of the aged SCN have revealed persistent cycling of clock gene expression but a breakdown of output signals^4,5^. However, whether age-related changes in systemic signaling impact tissue/organ clocks has yet to be elucidated. We demonstrate here that while the core clock continues to cycle in cultured fibroblasts synchronized with serum from old volunteers, the circadian transcriptome is different from that seen in cells treated with serum from young individuals. Through this analysis of the role of serum in age-induced changes in circadian rhythms, we suggest a potential mechanism for observations where specific genes showed a loss, gain, or maintenance of rhythmicity with age^19–23^. This phenomenon, known as circadian reprogramming, may illuminate which pathways are affected by or become more impactful for changes in cellular integrity through aging^24^.

In order to study the effect of circulating factors on age-related changes of the circadian transcriptome, we utilized the well-established serum starvation-serum addition protocol^7^ to synchronize cells in culture. Cultured fibroblasts from old and young subjects have robust clocks and respond similarly to synchronization with dexamethasone^9^; however, when synchronized with dexamethasone in a media containing old serum, they reportedly exhibited shortened circadian periods^9^. This effect was reversed by heat inactivating old serum. We did not observe a significant difference in period length between young and old serum synchronization in our *BMAL1-luciferase* experiment, perhaps because dexamethasone was not utilized. Our use of serum to synchronize allowed us to more closely simulate *in vivo* conditions where blood is an important mode by which central clocks can entrain peripheral clocks to maintain synchrony with the day:night cycle. Previously, serum factors were shown to activate the immediate early transcription factor, SRF, in a diurnal manner to confer time of day signals to both cells in culture and to mouse liver^7^. We show here that effects of age are also mediated by serum.

We find that while the number of rhythmic transcripts (weighted BIC>0.75) in the young serum condition (∼18%) is higher than the old serum condition (∼12%), both conditions demonstrate a much larger number of cycling transcripts in these cultured fibroblasts than in other cell culture studies^25–28^. In mammals, up to 20% of transcripts can cycle in a given tissue^29,30^, and the low number of cycling transcripts in culture has been cited as a major limitation of the culture model^27^. While our analysis used a normal human fibroblast cell line (BJ-5TA) we had higher statistical power, given that we had four different biological replicates at every 2-hour timepoint. However, a major contributing factor to the high rhythmicity might be the use of human serum as the synchronization signal. In this way, our model of mimicking signaling to peripheral tissues by using serum directly from humans may more accurately recapitulate the human condition. In this study we focused only on the effects of serum synchronization on fibroblasts, but it is likely that factors circulating in serum act on several tissues, and so their effects are relatively broad. However, we acknowledge that in order to support this claim future studies should investigate other peripheral tissue cell types. Additionally, in the future we intend to analyze the serum using a combination of fractionation and either proteomics or metabolomics to identify relevant factors for the regulation of peripheral rhythms.

Interestingly, many of the genes in the circadian transcriptome that exhibit age-related changes are independently implicated in aging. For instance, some of the genes that lose rhythmicity in the aged condition are involved in oxidative phosphorylation and mitochondrial function, both of which decrease with age^31^. Previous work in the field demonstrates that synchronization of the circadian clock in culture results in cycling of mitochondrial respiratory activity^32,33^ further underscoring the different effects of old serum, which does not support oscillations of oxidative phosphorylation associated transcripts. Age-dependent decrease in oxidative phosphorylation and increase in mitochondrial dysfunction^31^ is seen also in aged fibroblasts^34^ and contributes to age-related diseases^35^. We suggest that the age-related inefficiency of oxidative phosphorylation is conferred by serum signals to the cells such that oxidative phosphorylation cycles are mitigated. On the other hand, loss of cycling could contribute to impairments in mitochondrial function with age.

Both pathway analyses utilized here identified increased MESORs of steroid biosynthesis components in the aged sera condition. In particular, the genes identified in this pathway are associated with cholesterol biosynthesis. Since elderly individuals typically have lower levels of cholesterol biosynthesis and higher levels of circulating cholesterol^36^, we were surprised to see increased expression of biosynthesis transcripts. Perhaps the cholesterol synthesis related RNA levels are high as a response to low levels of cholesterol synthesis proteins within the cell. Previous studies in cultured hepatocytes demonstrated that increased reactive oxygen species resulted in higher levels of transcripts associated with cholesterol biosynthesis^37^. Given that deficits in oxidative phosphorylation are already implicated in these findings, it is possible that oxidative stress plays a role in the increase of cholesterol biosynthesis transcripts.

Although our findings are largely supported by the aging literature, the relatively small sample size of our study necessitates follow up studies to control for individual differences between subjects. We observed variations in luminescence and transcript traces across individuals and while we did not see changes that could account for the overall significant differences in transcript cycling between young and old subjects, we cannot exclude the possibility that factors other than aging contribute to these data.

Together, these findings indicate that at least some of the age-related changes in the cultured fibroblast circadian transcriptome are derived from signals circulating in the serum and not the age of the cells. This has profound implications for understanding and treating circadian disruption with age, and could also be relevant for other age-related pathology, given established links between circadian disruption and diseases of aging. Notably, many of the genes whose cycling is affected by old serum contribute to age-associated disorders.

## MATERIALS AND METHODS

### Clinical research study

This clinical research study enrolled apparent healthy participants from the volunteer pool maintained by the Institute for Translational Medicine and Therapeutics (ITMAT), University of Pennsylvania. The Institutional Review Board of the University of Pennsylvania (Federal wide Assurance FWA00004028; IRB Registration: IORG0000029) approved the clinical study protocol (Penn IRB#832866). The study was registered on ClinicalTrials.gov with identifier NCT04086589. After obtaining informed consent from all volunteers, study assessments were conducted in the Center for Human Phenomic Science (Penn CHPS#3002) in accordance with relevant GCP guidelines and regulations. We originally intended on recruiting n=20 per group; however, patient recruitment was halted due to the COVID-19 pandemic. Participants met criteria for inclusion (in general good health, either 70-85 years of age for the elderly cohort or 20-35 years of age for the young cohort, and a wrist-actigraphy-based average TST (total sleep time) ≥ 6 hours per night occurring between 22:00 – 08:00). Participants were excluded due to pregnancy or nursing, shift work (defined as recurring work between 22:00-05:00), a history of clinically significant obstructive sleep apnea, transmeridian travel across ≥2 time zones in the two weeks prior to study assessments and one week after, > 2 drinks of alcohol per day, and use of illicit drugs. The subject-specific midsleep preferences were quantified using the Munich ChronoType Questionnaire (MCTQ)^38^. Blood collections were done from the median cubital vein via venipuncture using a 22 G butterfly needle (BD, Franklin, Lakes, NJ, USA). All blood draws occurred at 14:00, the same time as Pagani et al.^9^. One subject returned for a repeat clinical assessment including biosampling to provide additional sample.

### Acquisition of accelerometry data streams

Participants wore a triaxial actigraph device (wGT3X-BT, ActiGraph, Pensacola, FL) on the non-dominant wrist. The devices were initialized using the following parameters: start date and time were synchronized with atomic server time without pre-defined stop date/time, at 60 Hz sampling rate for the three accelerometer axes, enabled for delay modus, steps, lux, inclinometer, and sleep while active. Raw data were downloaded from the device in AGD and GT3X file format in one second epochs using ActiLife software (version 6, ActiGraph, Pensacola, FL) and submitted for further analyses. For visualization and cosinor analysis, actigraphy data were aggregated into 1-minute intervals by summing ActiGraph counts across each minute. Cosinor analyses of the data were adapted from the single component cosinor analysis reviewed by Cornelissen^39^, as well as the cosine fit described by Refinetti et al. ^40^. Briefly, the measurement times for the actigraphy data were recalculated as hours since midnight on the first day of measurement, within each participant’s data. For each actigraphy variable and each participant, the lm() function in R (v4.2.0) was used to perform a cosinor fit with a fixed 24-hour period. The two cosinor coefficients from these fits were used to calculate participant-level amplitudes and circadian phases, while the intercepts provided MESOR estimates. The two-sided Wilcoxon rank sum exact test, as implemented by R’s wilcox.test() function, was used to test for significant differences in amplitude and MESOR between the age groups. A two-sample Kuiper’s test, as implemented by the kuiper_test(nboots = 10000) function from the twosamples R package (v2.0.0), was used to test for significant differences in circadian phase between the age groups.

### Acquisition of EKG data

The Zephyr BioPatch devices (Zephyr Technology, Annapolis, MD) were deployed as previously established^41^. All subjects included in this analysis wore the BioPatch for at least 24hrs. EKG recordings were analyzed by Kubios HRV Premium (ver. 3.5.0, Kubios Team, Kuopio, Finland) to report time-of-day-specific measures of cardiovascular function consisting of heart rate, RR intervals, sympathetic and parasympathetic nervous activity (SNS and PNS). Preprocessing in Kubios was set to automatic beat correction to remove artifacts and Smoothn priors for detrending. Cosinor fits and tests for differences in circadian parameters between age groups were performed as described for the actigraphy data. The circadian parameters MESOR and amplitude were tested for significant differences between the age groups by Wilcoxon rank sum exact test (two-sided), while phase was tested by Kuiper’s two-sample test to account for the circular measurement.

### Cortisol Measurements

Cortisol was measured in human serum by coated tube RIA (MP Bio, Solon OH) in duplicate. Tubes were counted on Perkin Elmer gamma counter and data reduced by STATLia software.

### Generation of stable BJ-5TA cell line expressing *BMAL1-dLuc-GFP*^10^

Virus was generated and cells were infected as we’ve previously described^42^. Briefly, LentiX 293T cells (Clonetech) were transfected with Lipofectamine 3000 PLUS (Life Tech) using manufacturer instructions. The transfection included 18ug of DNA per reaction and the plasmid (BMAL1-dLuc-GFP) to packaging vector (DVPR, Addgene) to envelope (VSV-G, Addgene) ratio was 10:1:0.5. Media was changed 24hrs post-transfection after the cells were checked using a fluorescence scope to make sure cells were >50% GFP positive. Supernatant with the virus was collected at 48hrs and 72hrs post transfection and spun down at 3000 RPM for five minutes (to eliminate any cells/debris). BJ-5TA cells were infected with fresh virus upon virus collection. Polybrene (Sigma-Aldrich, 10 mg/mL) was also added to aid with infection. Transduced BJ-5TA cells (>2000 cells) were sorted (FACSMelody, BD Biosciences) for high GFP expression. Once the cell line was established, blasticidin was added to the culture at 2ug/ml. Due to fragility of the cell line in the presence of antibiotic, BJ-5TA *Bmal1-luciferase* cells with differing expression levels of GFP were sorted. The line with highest stable luminescence oscillation was used for all experiments reported here.

### Serum entrainment and bioluminescent recording

BJ-5TA *BMAL1-dLuc-GFP* cells in 24-well plates (∼confluent) were washed (2x) with DPBS and given serum free media for 24hrs. After the starvation, cells were given media with 10% human serum and 200uM beetle luciferin potassium salt. Each well of cells was given media with the serum from a single patient. The cells were continuously monitored by LumiCycle luminometer (Actimetrics) for 4.5 days from the point of serum addition. LumiCycle raw data was exported using LumiCycle software (Actimentrics). The data were analyzed by BioDare2 (biodare2.ed.ac.uk^43^) using FFT NLLS with baseline detrending. Any replicates that were not cycling or had a period outside of the 20-28hr range was excluded from analysis. Both the serum free media and serum added media used the recipe from^10^ at pH 7.4; however, the serum free media had no serum and the serum media had 10% human serum instead of FBS.

### Sample preparation, RNA extraction, and RNA sequencing

BJ-5TA *BMAL1-dLuc-GFP* cells in 24-well plates (∼confluent) were washed (2x) with DPBS and given serum free media (DMEM with Penn Strep) for 24hrs. After 24hrs cells were given media with human serum (DMEM, Penn Strep, 10% human serum). Serum starvation was staggered every 12 hours over two days to allow for samples to be collected on the same day. Upon sample collection wells were place on ice and rinsed with cold DPBS and then put in cold RLT buffer with 2-Mercaptoethanol (10µL/mL). Samples were frozen at -80 overnight and sent to Admera for RNA extraction and sequencing. Total RNA was extracted with RNeay mini kit (Qiagen). Isolated RNA sample quality was assessed by High Sensitivity RNA Tapestation (Agilent Technologies Inc., California, USA) and quantified by Qubit 2.0 RNA HS assay (ThermoFisher, Massachusetts, USA). Paramagnetic beads coupled with oligo d(T)25 are combined with total RNA to isolate poly(A)+ transcripts based on NEBNext® Poly(A) mRNA Magnetic Isolation Module manual (New England BioLabs Inc., Massachusetts, USA). Prior to first strand synthesis, samples are randomly primed (5 ’ d(N6) 3 ’ [N=A,C,G,T]) and fragmented based on manufacturer’s recommendations. The first strand is synthesized with the Protoscript II Reverse Transcriptase with a longer extension period, approximately 40 minutes at 42°C. All remaining steps for library construction were used according to the NEBNext® UltraTM II Non-Directional RNA Library Prep Kit for Illumina® (New England BioLabs Inc., Massachusetts, USA). Final libraries quantity was assessed by Qubit 2.0 (ThermoFisher, Massachusetts, USA) and quality was assessed by TapeStation D1000 ScreenTape (Agilent Technologies Inc., California, USA). Final library size was about 430bp with an insert size of about 300bp. Illumina® 8-nt dual-indices were used. Equimolar pooling of libraries was performed based on QC values and sequenced on an Illumina® NovaSeq S4 Illumina, California, USA) with a read length configuration of 150 PE for 40 M PE reads per sample (20 M in each direction).

### RNA-seq and statistical analysis

Raw RNA-seq reads were aligned to the GRCh38 build of the human genome by STAR version 2.7.10a^44^. The dataset contained an average of 19,265,220 paired-end non-stranded 150 bp reads mapping uniquely to genes, per sample. Data were normalized and quantified at both gene and exon-intron level, using a downsampling strategy implemented in PORT (Pipeline Of RNA-seq Transformations, available at https://github.com/itmat/Normalization), version 0.8.5f-beta_hotfix1. Both STAR and PORT were provided with gene models from release 106 of the Ensembl annotation^45^.

MESOR, amplitude, and phase estimates, as well as p-values for the difference in MESOR, amplitude, and phase, were calculated with CircaCompare^13^, version 0.1.1. Only rhythmic genes were taken into consideration in pathway and other analyses involving MESORs and phases. The criterion for rhythmicity was either based on (BH adjusted) p-values reported by CircaCompare, or weighted BIC values, obtained by an approach similar to dryR^46^. In the latter approach we fitted four models of rhythmicity, one modeling gene expression that is rhythmic in both cells treated with young or old sera, another modeling gene expression rhythmic in neither cells treated with young nor old sera, and two modeling gene expression rhythmic in either young or old sera treated cells respectively. The BIC values were calculated for the four models and were weighted to obtain numbers between 0 and 1. All methods were provided with log_10_ (1 + PORT normalized count) values and were run on R, version 4.1.2, accessed through Python, version 3.9.9, via rpy2, an interface to R running embedded in a Python process, https://rpy2.github.io/, version 3.4.5. Additionally, we used Nitecap^47^ to visualize and explore circadian profiles of gene expression.

### STRING pathway analysis

We performed STRING^14^ pathway analyses using STRING API version 11.5. Enrichment analyses were performed on the sets of genes rhythmic in both groups (weighted BIC > 0.75) with observed decrease in MESOR, increase in MESOR, advance in phase, and delay in phase according to CircaCompare q < 0.05 criterion. We also performed analyses on the sets consisting of genes rhythmic only in young group (weighted BIC > 0.75), only in old group (weighted BIC > 0.75), and the set of genes rhythmic in both groups (weighted BIC > 0.75). Finally, two additional analyses were performed, on the sets of genes with observed decrease and increase of amplitude (CircaCompare q < 0.05).

### Ingenuity Pathway Analysis

QIAGEN IPA^15^ was used to identify pathways enriched in various subsets of cycling genes. The following subsets of genes were identified using a combination of CircaCompare stats and BIC cutoffs: (1) Decreased MESOR in old sera (929 genes) – CircaCompare rhythmic q < 0.05 in old, CircaCompare rhythmic q < 0.05 in young, CircaCompare MESOR difference q < 0.05, MESOR difference (old – young) < 0. (2) Increased MESOR in old sera (515 genes) – same selection criteria as ‘Decreased MESOR in old sera,’ except MESOR difference (old – young) > 0. (3) Phase advance in old sera (148 genes) - CircaCompare rhythmic q < 0.05 in old, CircaCompare rhythmic q < 0.05 in young, CircaCompare Phase difference q < 0.05, phase difference (old – young) < 0. (4) Phase delay in old sera (156 genes) – same selection criteria as ‘Phase advance in old sera,’ except phase difference (old – young) > 0. (5) Loss of rhythmicity in old sera (1519 genes) – weighted BIC > 0.75 for ‘Rhythmic in Young but not Old’ model. (6) Gain of rhythmicity in old sera (637 genes) – weighted BIC > 0.75 for ‘Rhythmic in Old but not Young’ model. (7) Rhythmic in old and young sera (1209 genes) – weighted BIC > 0.75 for ‘Rhythmic in both Old and Young’ model. Each of these gene subsets were processed separately with IPA’s core analysis, using default parameters.

For visualization via heatmap, we perform three rounds of normalization within each gene. First, we mean-normalize the read counts within each age group and serum treatment group (A, B, C, D). This is to account for baseline differences between sera collected from the different subjects. Second, we collapse replicates at each timepoint by calculating their means. Third, we calculate Z-Scores across all timepoints, within each age group. Note, these normalization procedures are to aid with visualization of the data and were not used as part of the statistical analyses.

### Epigenetic Landscape in Silico deletion (LISA)

LISA analysis^48^ was used to perform transcription factor binding analysis. LISA results were filtered by fibroblast. q<0.05 cut off was used within each of the 5 groups. Venn diagram was generated with https://bioinformatics.psb.ugent.be/webtools/Venn/

### Data and materials availability

Sequencing data are deposited in Gene Expression Omnibus (NCBI) under accession number GSE270290. All additional data files are available upon request.

## Acknowledgments

We wish to extend our sincere gratitude to the study volunteers. Ms. LaVenia Banas provided excellent logistical study support. We thank Dr. Andrew Liu for the BMAL1-luc plasmid. We thank the RIA Biomarker Core of the Penn Diabetes Research Center, P30-DK19525 for cortisol measurements. We thank Sara Bernardez-Noya and Rebecca Moore for input on analysis and data presentation. The project described was supported by the National Center for Research Resources and the National Center for Advancing Translational Sciences, National Institutes of Health, through Grant 5UL1TR001878 (A.S.) and National Heart, Blood, and Lung Institutes of Health, through Grant R00HL147212 (S.L.Z.). C.S. is the Robert L. McNeil Jr. Fellow in Translational Medicine and Therapeutics. A.S. is an investigator of the Howard Hughes Medical Institute. J.E.S. was supported by a training grant in Neuroscience (NIH T32-NS105607), an National Institutes of Health Diversity Supplement (NIH NS48471), and by a grant to the University of Pennsylvania from the Howard Hughes Medical Institute through the James H. Gilliam Fellowship for Advanced Study program. The funders had no role in study design, data collection and analysis, decision to publish, or preparation of the manuscript.

---

**Fig S1.**
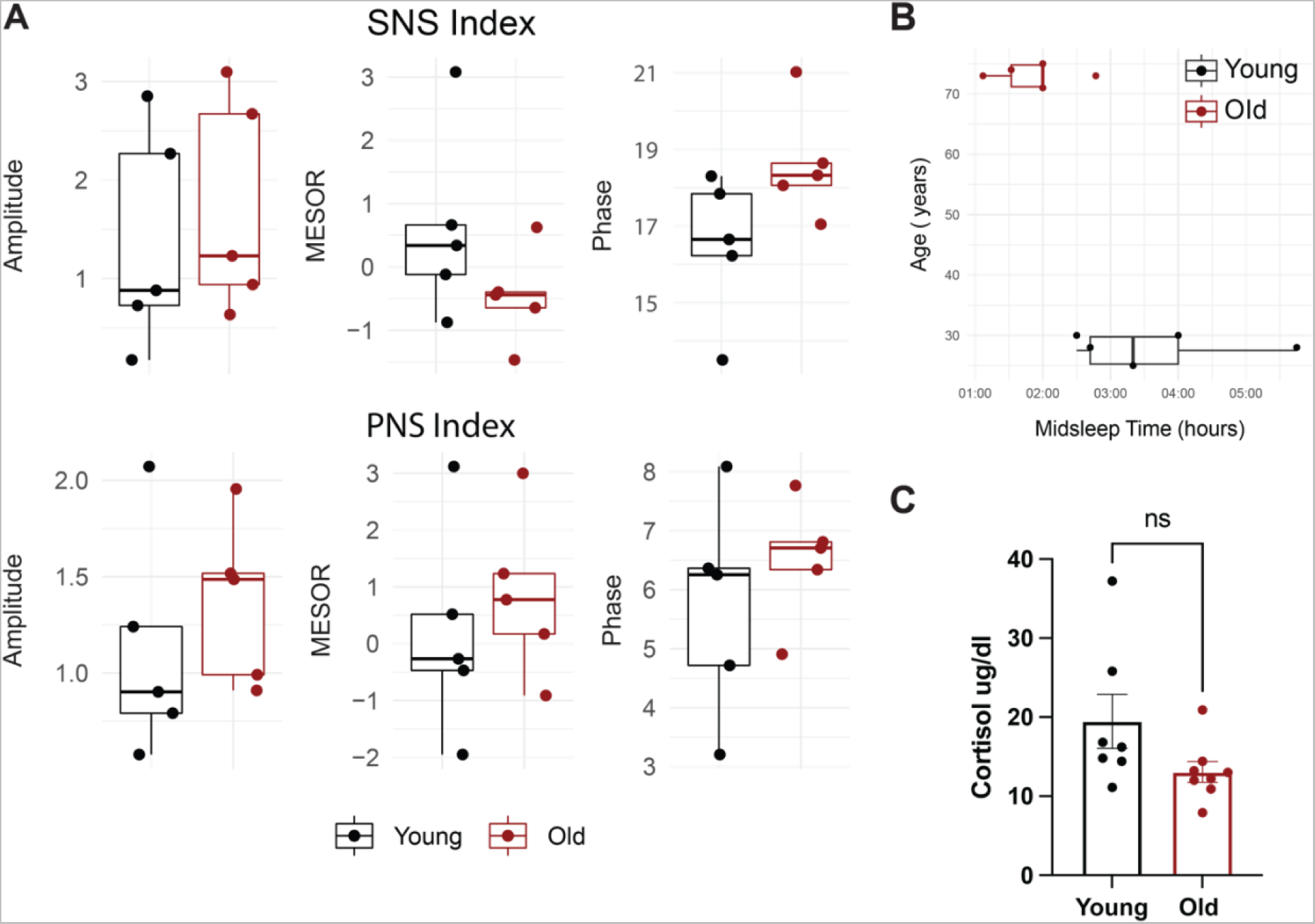
There are differences in midsleep time, but no significant differences in sympathetic, parapsympathetic nervous system indices, or cortisol levels between young and old individuals. Sympathetic nervous system (SNS, A, top) and parasympathetic nervous system (PNS, A, bottom) indices were measured using EKG. Amplitude, MESOR, and Phase were calculated from the resulting oscillations and the Median, quartiles, and SEMs are shown. Dots represent individuals. P>0.05. N=5 per group. MESOR and amplitude were tested using Wilcoxon rank sum exact test, while phase was tested by Kuiper’s two-sample test. Boxplot midlines correspond to median values, while the lower and upper hinges correspond to the first and third quartiles, respectively. Boxplot whiskers extend to the smallest/largest points within 1.5 * IQR (inter Quartile Range) of the lower/upper hinge (A). Boxplot distributions of midsleep times for subjects in the young (black) and old (red) cohorts. The midsleep time is advanced in the old subjects compared to young by Mann-Whitney U test (Wilcoxon rank sum test) p=0.036). We calculated midsleep times for each subject from their responses to the Munich Chronotype Questionnaire (MCTQ). N=5 per group. We were not able to calculate midsleep times for four subjects (two from the old cohort and two from the young cohort) because they used alarms to wake on their non-working days and one old subject because they exhibited a total sleep time less than 6 hours (B). Serum cortisol levels are not statistically different by unpaired t-test (C, N=7 young, 8 old).

**Figure S2.**
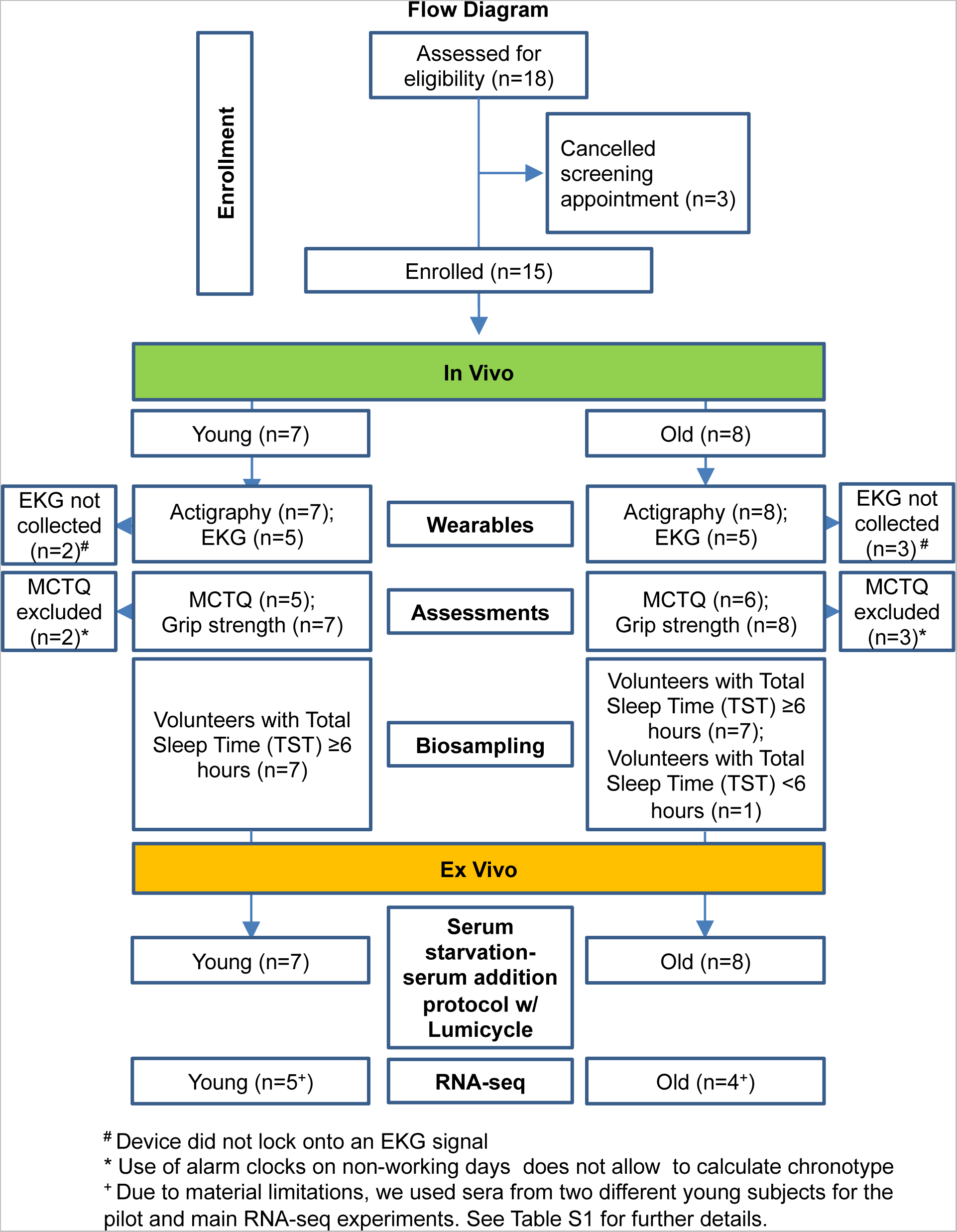
Subject Inclusion Flow Chart. A flow chart of the number of subjects included in each analysis present in this study.

**Fig S3:**
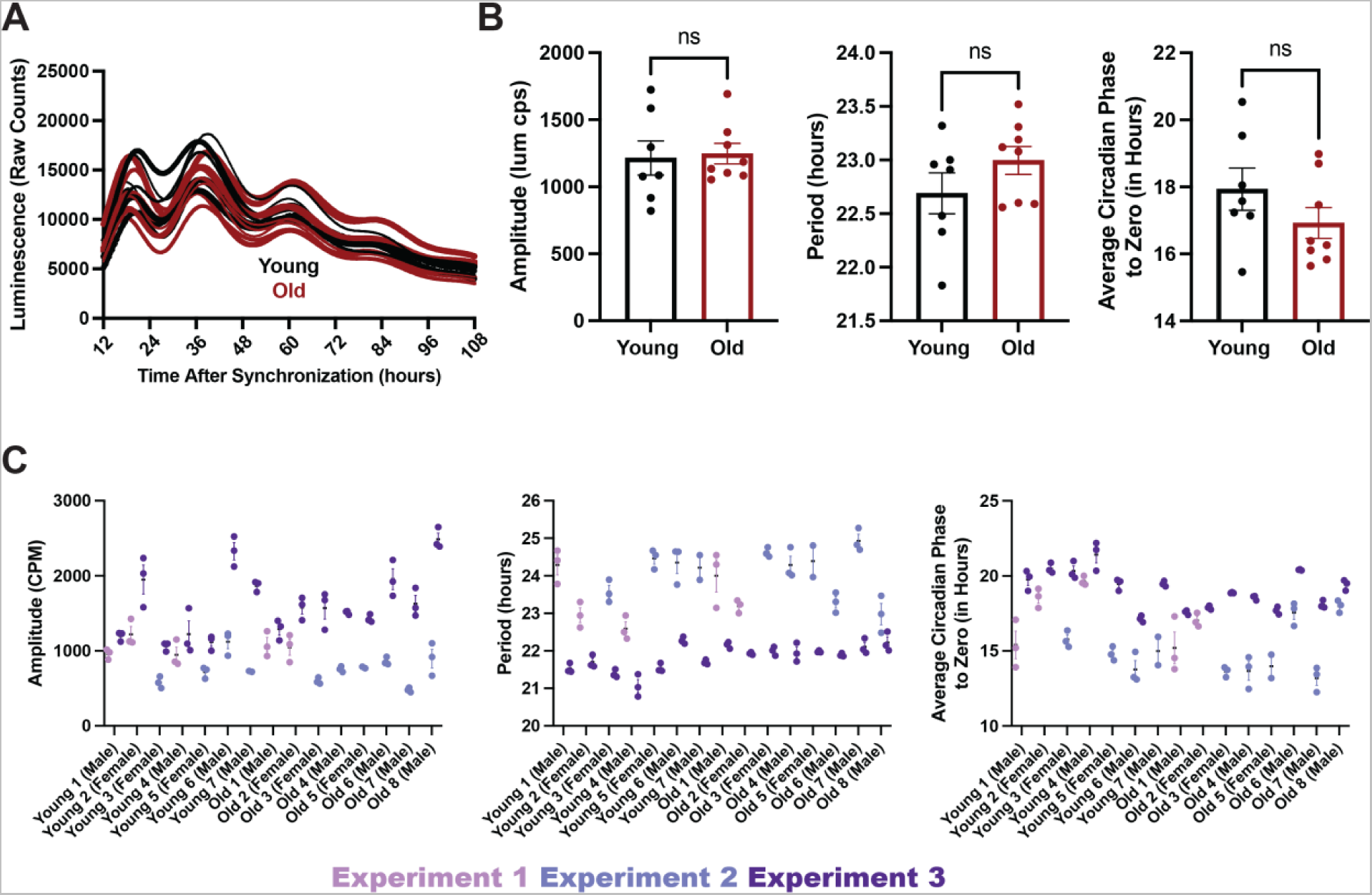
Young and old serum are equally effective at entraining cells in culture. **(A,B)** Cells synchronized with human serum from either young (N=7 subjects) or old (N=8 subjects) individuals did not show differences in amplitude (as measured in luminescence counts per second, bottom, left), period (bottom, middle), or phase (bottom, right). Line traces and data points represent average values for a specific patient’s serum over 2 experiments with 2-3 replicates per experiment. A visual representation of the individual replicates averaged in B **(C)** Summary statistics are displayed as mean +/- SEM. Means compared by unpaired t-test. Despite individual and run to run variability, cells synchronized with human serum from either young (N=7 subjects) or old (N=8 subjects) individuals showed similar amplitude (left), period (middle), and phase relative to synchronization time (right) of BMAL1-luciferase rhythms. Data points represent individual replicates within an experiment. Wells run in the same experiment are displayed in the same color. Each subject’s sample was run in 2-3 replicates.

**Fig S4.**
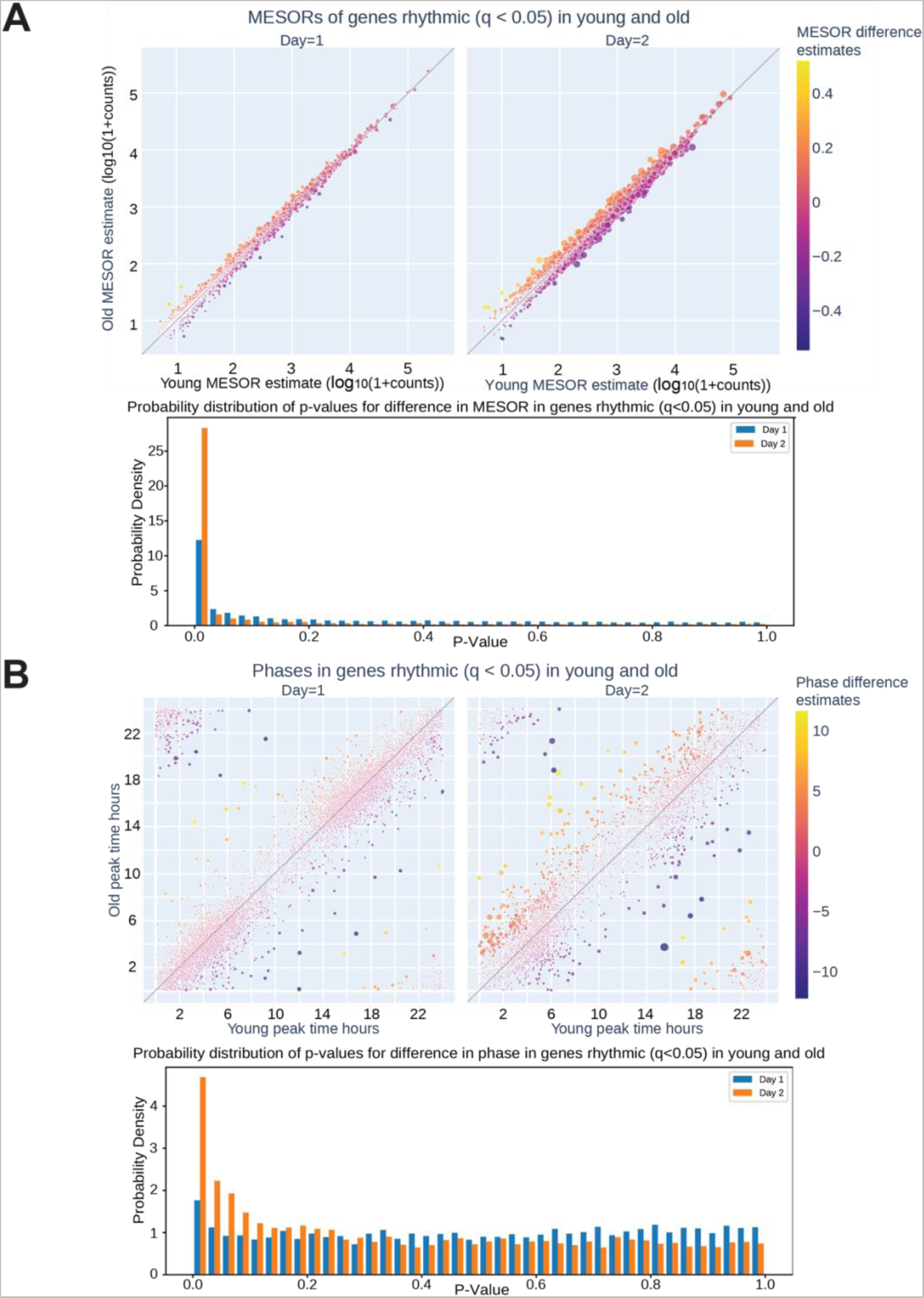
The circadian transcriptomes of cells synchronized with young or old sera deviate significantly on day two. On day two of serum entrainment, young and old transcriptional rhythms differentiated. CircaCompare analysis of RNA sequencing revealed that on Day 2 of serum entrainment analysis (36-58 hours after synchronization) the MESOR differences of cycling genes are larger between the cells entrained with young or old serum **(A**, top). Additionally, more genes were phase shifted in the old serum condition compared to young serum on Day 2 **(B**, top). The distribution of p-values shows an enrichment of low p-values on Day 2 for both MESOR and phase differences (A, bottom, B, bottom). The size of each circle in the scatter plots is proportional to -log_10_q, hence bigger circles correspond to smaller q-values for the difference between young and old for each metric.

**Figure S5.**
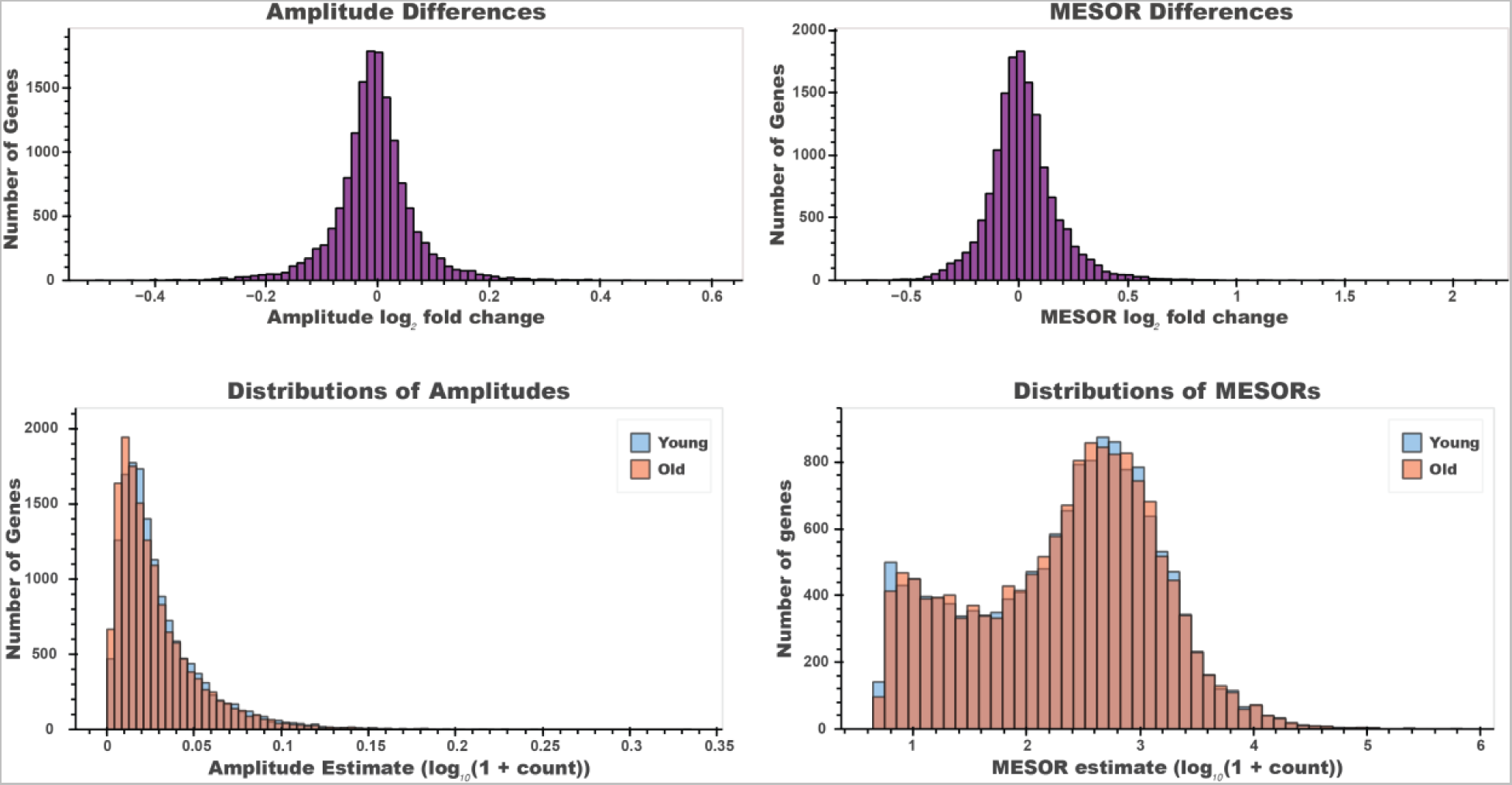
There is a small effect of age on the amplitude and MESOR of cycling mRNA between young and old serum samples. The difference (Top, left) between young and old mRNA amplitude values (bottom, left) are significantly different by Wilcoxon signed-rank test (p= 3.73e-39). The difference (Top, right) between young and old mRNA MESOR values (bottom, right) are significantly different by Wilcoxon signed-rank test (p= 6.94e-22). However, these small differences result in very low p-values due to the large number of genes inspected.

**Figure S6:**
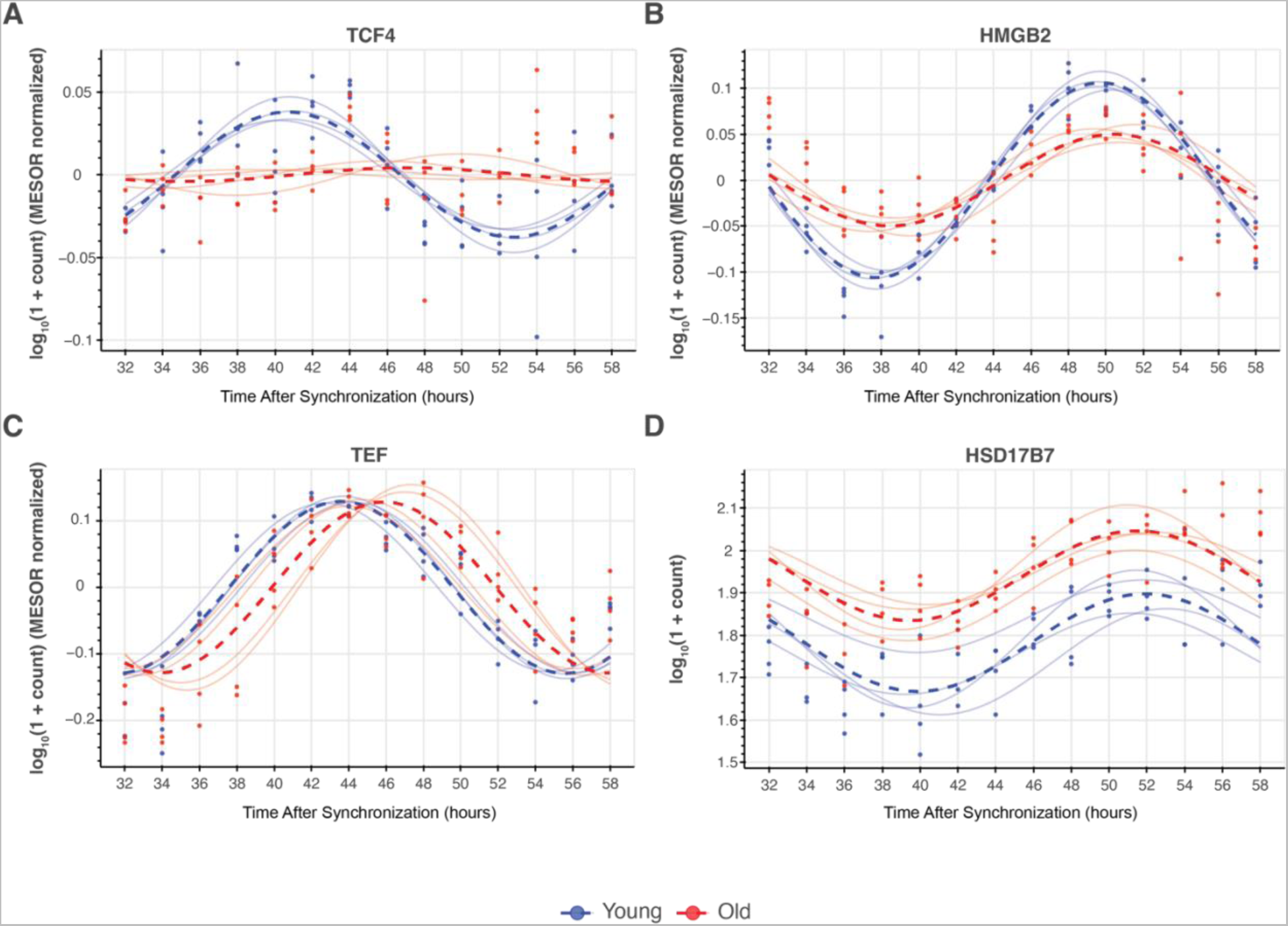
Sample traces of transcripts that cycle with young serum synchronization and are differentially affected by old serum synchronization. Example traces of mRNA transcripts that lose cycling (A, TCF4), decrease amplitude (B, HMGB2), phase shift (C, TEF), and increase MESOR (D, HSD17B7). To ease the comparison of amplitudes and phases in traces depicted in (A-C), subject-level differences in MESORs were removed by subtracting the measured gene expression of each subject at each timepoint by the MESOR of the subject’s gene expression oscillation pattern, making all subject-level traces oscillate around zero. Traces in (D) were not subjected to this transformation since (D) showcases differences in MESORs themselves. Transparent lines represent subject-level fits and dashed/opaque lines represent cohort-level fits.

**Fig S7:**
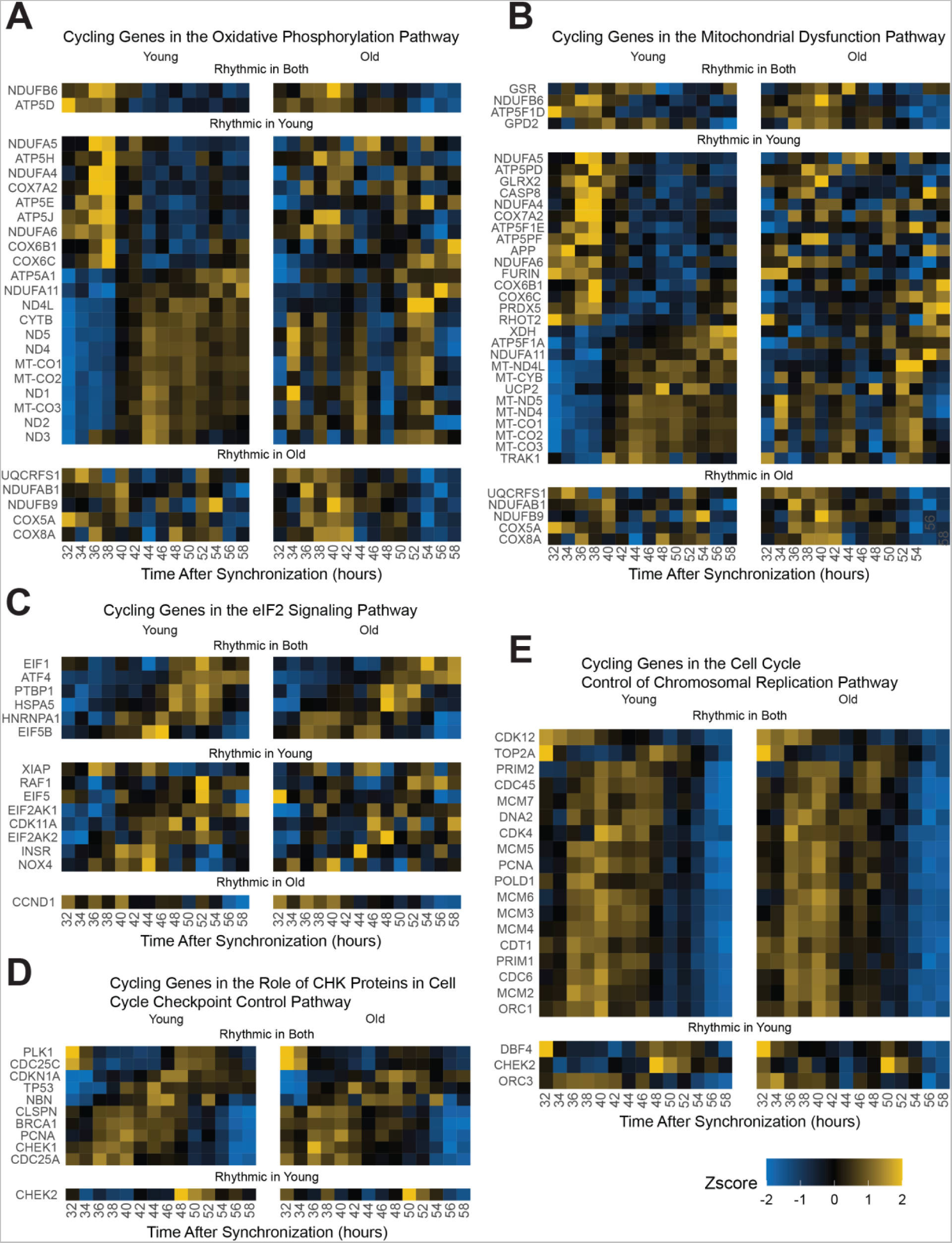
Oxidative phosphorylation genes and chromosomal replication genes are affected by entrainment with old serum. Additional pathway analysis employing IPA further emphasizes finding with STRING analysis. Genes identified by IPA analysis to be associated with oxidative phosphorylation/mitochondrial dysfunction and eIF2 genes lose rhythmicity with age (BIC >0.75) **(A, B, C)**. Cell cycle checkpoint control (CHK proteins) **(D)** and chromosomal replication genes **(E)** maintain their cycling in the aged condition, which is consistent with the continued division of cells.

**Fig S8:**
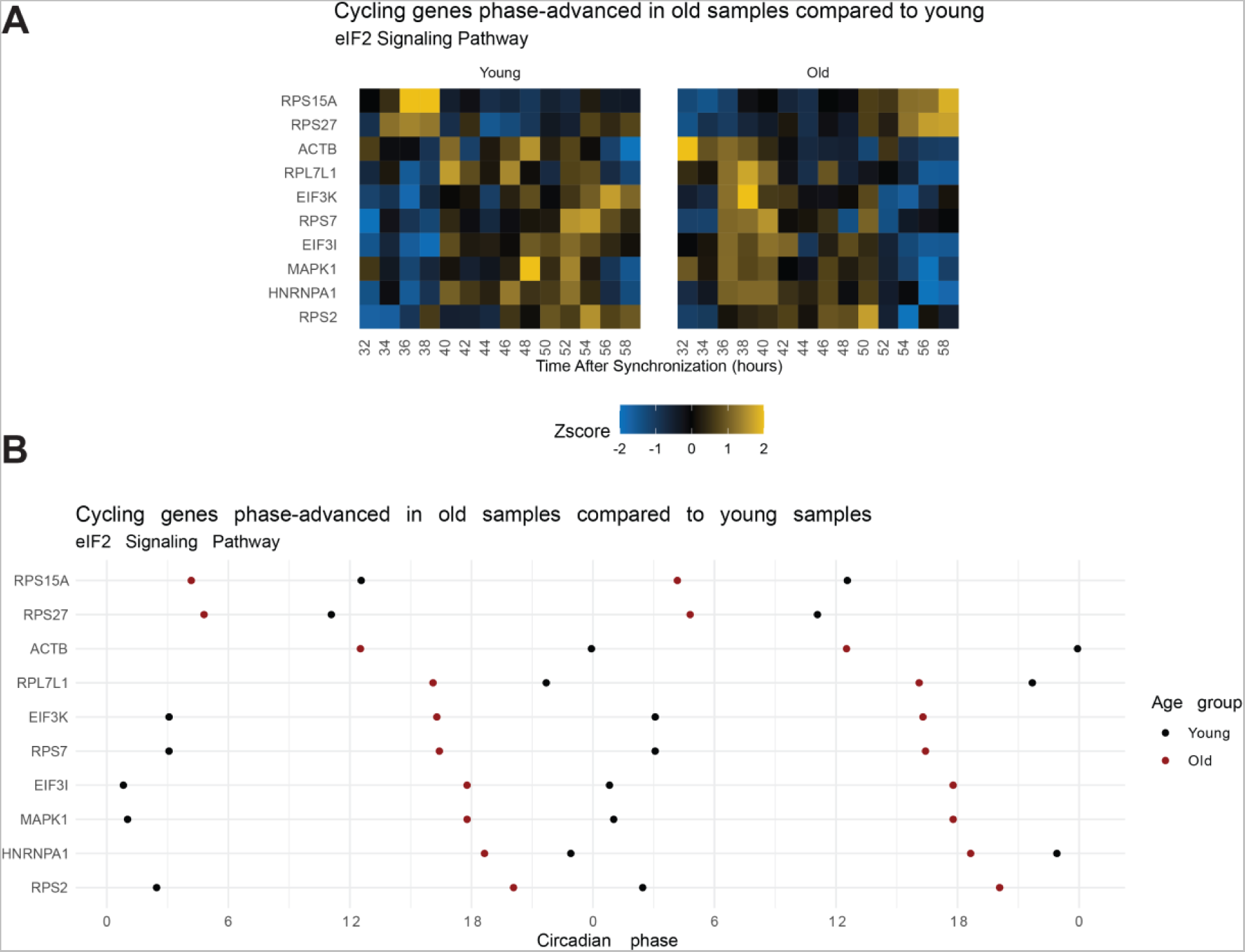
eIF2 signaling pathway genes that maintain cycling with age phase advance in the old serum condition. From the IPA analyses, the eIF2 signaling pathway was enriched (BIC >0.75) in two datasets: 1) genes cycling in the young sera and not the old and 2) cycling genes phase-advanced in the old sera (q<0.05). A significant number of genes that maintain rhythmicity in the old sera condition are phase advanced as visualized in the heat map (A) or plot of peak phase (B). Note, the data in the peak phase plot are double-plotted. This means the range of the x-axis is doubled and the datapoints between 0 and 24 are duplicated in the right half of the plot. Phase is a circular metric meaning a phase of 0 is equivalent to a phase of 24 (1 ⇔ 25, 2 ⇔ 26, etc). Double-plotting is used to visualize data with repeated patterns or when data straddle the 24-0 boundary. As the eIF2 pathway is involved in protein translation, this suggests that the timing of protein translation is phase advanced in the older serum and/or less rhythmic.

**Table S1.** Demographic information for subjects in the study.

| Age Group | Sex | Age (y) | BMI (kg/m<sup>2</sup>) | Total Sleep Time (TST) [h] | Actigraphy | MCTQ | Grip Strength | EKG | RNA-seq: Main | RNA-seq: Pilot (Fig S4) |
| --- | --- | --- | --- | --- | --- | --- | --- | --- | --- | --- |
| 20-35 | Female | 25 | 23.7 | 6.6 | Yes | Yes | Yes | No | No | No |
|  | Female | 28 | 21.2 | 6.9 | Yes | No | Yes | Yes | Yes | No |
|  | Male | 28 | 25 | 9.7 | Yes | Yes | Yes | Yes | Yes | No |
|  | Male | 28 | 25.2 | 7.4 | Yes | No | Yes | No | Yes | No |
|  | Male | 28 | 25.9 | 9.3 | Yes | Yes | Yes | Yes | No | No |
|  | Female | 30 | 28.7 | 7.4 | Yes | Yes | Yes | Yes | Yes | No |
|  | Male | 30 | 25.9 | 6.4 | Yes | Yes | Yes | Yes | No | Yes |
| **Young** | **Mean** | **28.1** | **25.1** | **7.7** | — | — | — | — | — | — |
|  | **SD** | **1.7** | **2.3** | **1.3** | — | — | — | — | — | — |
| 70-85 | Male | 70 | 26.8 | 6.7 | Yes | No | Yes | Yes | No | No |
|  | Male | 71 | 26.7 | 11.3 | Yes | Yes | Yes | No | Yes | No |
|  | Female | 73 | 32.4 | 7.7 | Yes | Yes | Yes | No | Yes | No |
|  | Male | 73 | 21.5 | 6.6 | Yes | Yes | Yes | Yes | No | No |
|  | Female | 74 | 26.4 | 7.5 | Yes | No | Yes | No | No | No |
|  | Male | 74 | 27.2 | 7 | Yes | Yes | Yes | Yes | Yes | Yes |
|  | Male | 75 | 28.8 | 7.8 | Yes | Yes | Yes | Yes | No | No |
|  | Female | 76 | 22.3 | 5.8 | Yes | Yes | Yes | Yes | Yes | No |
| **Old** | **Mean** | **73.3** | **26.5** | **7.6** | — | — | — | — | — | — |
|  | **SD** | **2** | **3.4** | **1.7** | — | — | — | — | — | — |

**Table S2.**
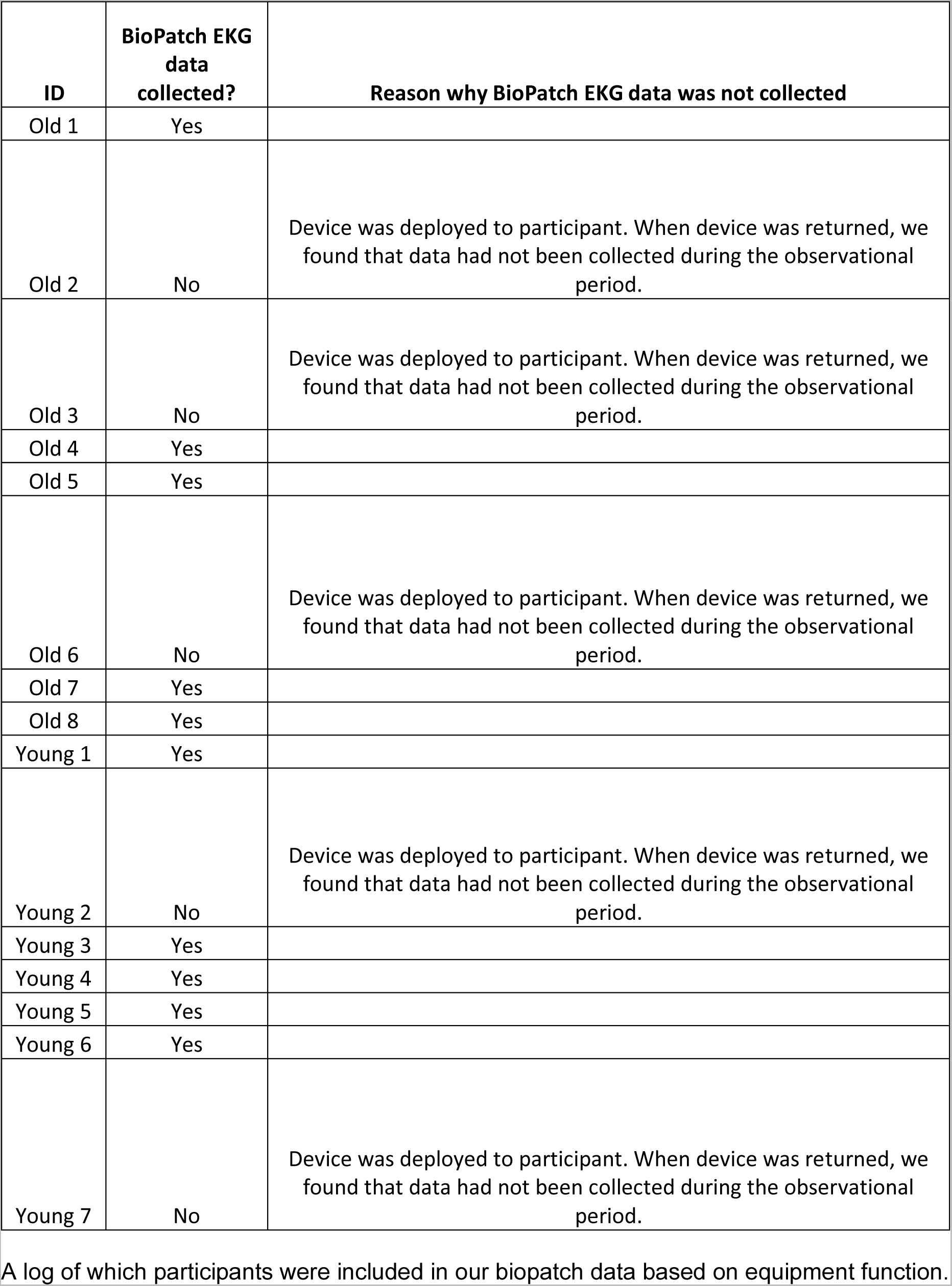
BioPatch EKG Data Inclusion Log.

**Table S3:**
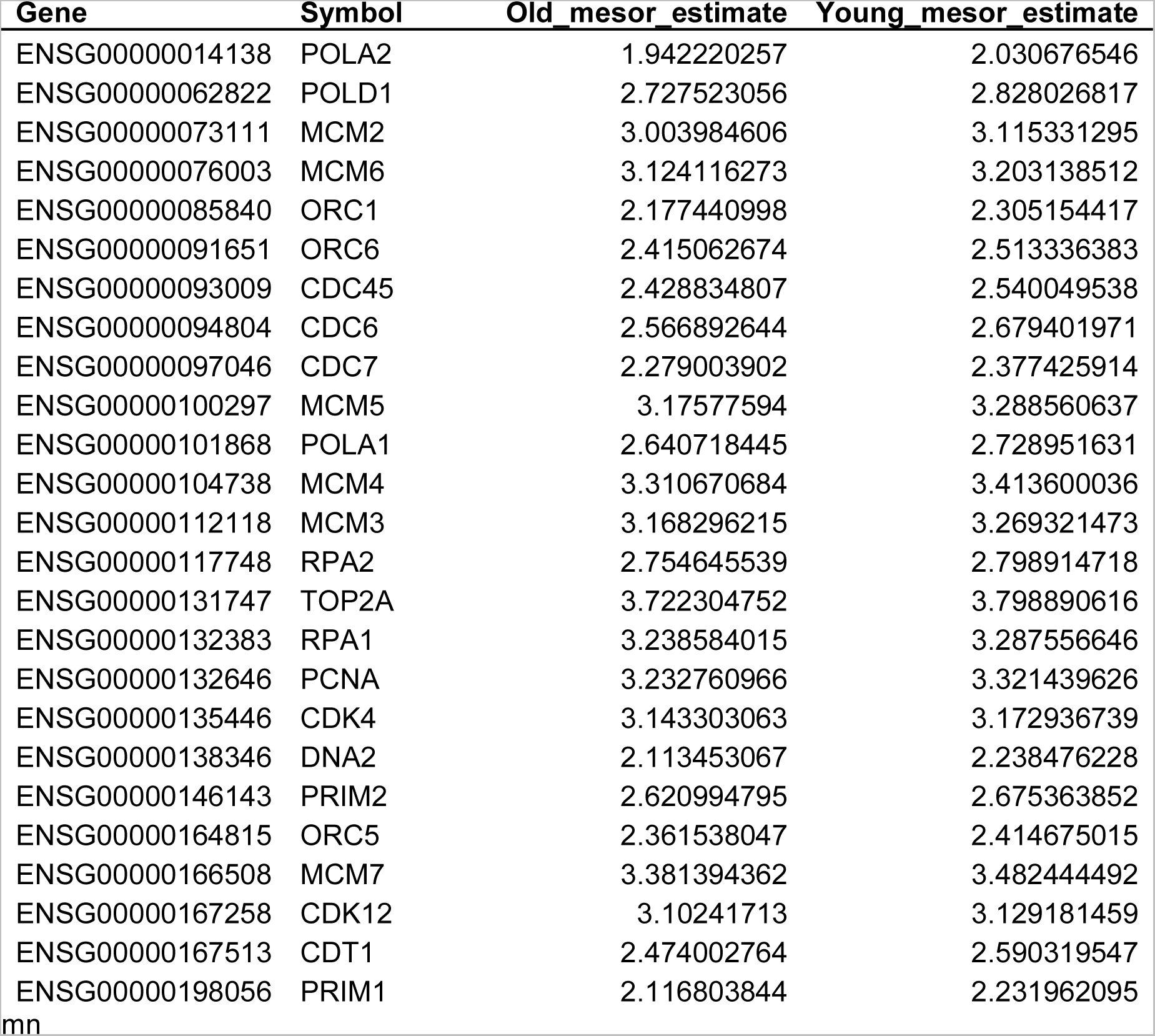
Genes involved in the IPA cell cycle of chromosome replication pathway show decreased MESOR with age. Table of genes that are rhythmic in both conditions with a decreased MESOR with old serum treatment compared to young serum treatment by CircaCompare. All genes are involved in the cell cycle/DNA replication pathway.

**Table S4:** Genes with increased MESOR in the IPA cholesterol biosynthesis pathway. Table of genes that are rhythmic in both conditions with a decreased MESOR with old serum treatment compared to young serum treatment by CircaCompare. All genes are involved in the cholesterol biosynthesis pathway.

| Gene | Symbol | Old_mesor_estimate | Young_mesor_estimate |
| --- | --- | --- | --- |
| ENSG00000001630 | CYP51A1 | 2.040377336 | 1.871207095 |
| ENSG00000079459 | FDFT1 | 3.330291993 | 3.19001773 |
| ENSG00000104549 | SQLE | 3.116822033 | 2.950386727 |
| ENSG00000110921 | MVK | 2.530172998 | 2.4121956 |
| ENSG00000112972 | HMGCS1 | 2.93111747 | 2.696086286 |
| ENSG00000113161 | HMGCR | 3.011982868 | 2.833316048 |
| ENSG00000116133 | DHCR24 | 3.506294106 | 3.343492054 |
| ENSG00000120437 | ACAT2 | 2.71791125 | 2.58109012 |
| ENSG00000132196 | HSD17B7 | 1.940436315 | 1.782521965 |
| ENSG00000147383 | NSDHL | 2.540302415 | 2.468852126 |
| ENSG00000160285 | LSS | 3.486802994 | 3.411431601 |
| ENSG00000160752 | FDPS | 3.143592562 | 3.081359846 |
| ENSG00000167508 | MVD | 2.819963909 | 2.68386468 |
| ENSG00000172893 | DHCR7 | 2.990620862 | 2.797567922 |

## References

1. Monk, T. H., Buysse, D. J., Reynolds, C. F. & Kupfer, D. J. Inducing jet lag in older people: Adjusting to a 6-hour phase advance in routine. Experimental Gerontology 28, 119–133 (1993).

2. Monk, T. H., Buysse, D. J., Carrier, J. & Kupfer, D. J. Inducing jet-lag in older people: Directional asymmetry. Journal of Sleep Research 9, 101–116 (2000).

3. Hood, S. & Amir, S. The aging clock: circadian rhythms and later life. Journal of Clinical Investigation 127, 437–446 (2017).

4. Farajnia, S. et al. Evidence for Neuronal Desynchrony in the Aged Suprachiasmatic Nucleus Clock. J. Neurosci. 32, 5891–5899 (2012).

5. Nakamura, T. J. et al. Age-Related Decline in Circadian Output. J Neurosci 31, 10201– 10205 (2011).

6. Yamazaki, S. et al. Effects of aging on central and peripheral mammalian clocks. Proc Natl Acad Sci U S A 99, 10801–10806 (2002).

7. Gerber, A. et al. Blood-Borne Circadian Signal Stimulates Daily Oscillations in Actin Dynamics and SRF Activity. Cell 152, 492–503 (2013).

8. Katsimpardi, L. et al. Vascular and Neurogenic Rejuvenation of the Aging Mouse Brain by Young Systemic Factors. Science 344, 630–634 (2014).

9. Pagani, L. et al. Serum factors in older individuals change cellular clock properties. Proceedings of the National Academy of Sciences of the United States of America (2011) doi:10.1073/pnas.1008882108.

10. Ramanathan, C., Khan, S. K., Kathale, N. D., Xu, H. & Liu, A. C. Monitoring Cell-autonomous Circadian Clock Rhythms of Gene Expression Using Luciferase Bioluminescence Reporters. JoVE (Journal of Visualized Experiments) e4234 (2012) doi:10.3791/4234.

11. Balsalobre, A., Damiola, F. & Schibler, U. A Serum Shock Induces Circadian Gene Expression in Mammalian Tissue Culture Cells. Cell 93, 929–937 (1998).

12. Al-Romaih, K. et al. Chromosomal instability in osteosarcoma and its association with centrosome abnormalities. Cancer Genetics and Cytogenetics 144, 91–99 (2003).

13. Parsons, R., Parsons, R., Garner, N., Oster, H. & Rawashdeh, O. CircaCompare: a method to estimate and statistically support differences in mesor, amplitude and phase, between circadian rhythms. Bioinformatics 36, 1208–1212 (2020).

14. Szklarczyk, D. et al. STRING v11: protein-protein association networks with increased coverage, supporting functional discovery in genome-wide experimental datasets. Nucleic Acids Res 47, D607–D613 (2019).

15. Krämer, A., Green, J., Pollard, J. & Tugendreich, S. Causal analysis approaches in Ingenuity Pathway Analysis. Bioinformatics 30, 523–530 (2014).

16. Holubiec, M. I., Gellert, M. & Hanschmann, E. M. Redox signaling and metabolism in Alzheimer’s disease. Front Aging Neurosci 14, 1003721 (2022).

17. Sehar, U., Rawat, P., Reddy, A. P., Kopel, J. & Reddy, P. H. Amyloid Beta in Aging and Alzheimer’s Disease. Int J Mol Sci 23, 12924 (2022).

18. Raulin, A.-C. et al. ApoE in Alzheimer’s disease: pathophysiology and therapeutic strategies. Mol Neurodegener 17, 72 (2022).

19. Chen, C.-Y. et al. Effects of aging on circadian patterns of gene expression in the human prefrontal cortex. Proc Natl Acad Sci U S A 113, 206–211 (2016).

20. Kuintzle, R. C. et al. Circadian deep sequencing reveals stress-response genes that adopt robust rhythmic expression during aging. Nat Commun 8, 14529 (2017).

21. Sato, S. et al. Circadian Reprogramming in the Liver Identifies Metabolic Pathways of Aging. Cell 170, 664–677.e11 (2017).

22. Solanas, G. et al. Aged Stem Cells Reprogram Their Daily Rhythmic Functions to Adapt to Stress. Cell 170, 678–692.e20 (2017).

23. Blacher, E. et al. Aging disrupts circadian gene regulation and function in macrophages. Nat Immunol 23, 229–236 (2022).

24. Wolff, C. A. et al. Defining the age-dependent and tissue-specific circadian transcriptome in male mice. Cell Reports 42, 111982 (2023).

25. Duffield, G. E. et al. Circadian programs of transcriptional activation, signaling, and protein turnover revealed by microarray analysis of mammalian cells. Curr Biol 12, 551–557 (2002).

26. Grundschober, C. et al. Circadian regulation of diverse gene products revealed by mRNA expression profiling of synchronized fibroblasts. J Biol Chem 276, 46751–46758 (2001).

27. Hughes, M. E. et al. Harmonics of Circadian Gene Transcription in Mammals. PLOS Genetics 5, e1000442 (2009).

28. Jang, C., Lahens, N. F., Hogenesch, J. B. & Sehgal, A. Ribosome profiling reveals an important role for translational control in circadian gene expression. Genome Res 25, 1836– 1847 (2015).

29. Patel, V. R., Eckel-Mahan, K., Sassone-Corsi, P. & Baldi, P. CircadiOmics: integrating circadian genomics, transcriptomics, proteomics and metabolomics. Nat Methods 9, 772– 773 (2012).

30. Zhang, R., Lahens, N. F., Ballance, H. I., Hughes, M. E. & Hogenesch, J. B. A circadian gene expression atlas in mammals: Implications for biology and medicine. Proc Natl Acad Sci U S A 111, 16219–16224 (2014).

31. Lesnefsky, E. J. & Hoppel, C. L. Oxidative phosphorylation and aging. Ageing Research Reviews 5, 402–433 (2006).

32. Cela, O. et al. Clock genes-dependent acetylation of complex I sets rhythmic activity of mitochondrial OxPhos. Biochimica et Biophysica Acta (BBA) - Molecular Cell Research 1863, 596–606 (2016).

33. Scrima, R. et al. Mitochondrial calcium drives clock gene-dependent activation of pyruvate dehydrogenase and of oxidative phosphorylation. Biochimica et Biophysica Acta (BBA) - Molecular Cell Research 1867, 118815 (2020).

34. Greco, M. et al. Marked aging-related decline in efficiency of oxidative phosphorylation in human skin fibroblasts. The FASEB Journal 17, 1706–1708 (2003).

35. Federico, A. et al. Mitochondria, oxidative stress and neurodegeneration. Journal of the Neurological Sciences 322, 254–262 (2012).

36. Bertolotti, M. et al. Age-associated alterations in cholesterol homeostasis: evidence from a cross-sectional study in a Northern Italy population. Clin Interv Aging 9, 425–432 (2014).

37. Seo, E., Kang, H., Choi, H., Choi, W. & Jun, H.-S. Reactive oxygen species-induced changes in glucose and lipid metabolism contribute to the accumulation of cholesterol in the liver during aging. Aging Cell 18, e12895 (2019).

38. Roenneberg, T., Wirz-Justice, A. & Merrow, M. Life between Clocks: Daily Temporal Patterns of Human Chronotypes. J Biol Rhythms 18, 80–90 (2003).

39. Cornelissen, G. Cosinor-based rhythmometry. Theor Biol Med Model 11, 16 (2014).

40. Refinetti, R., Lissen, G. C. & Halberg, F. Procedures for numerical analysis of circadian rhythms. Biol Rhythm Res 38, 275–325 (2007).

41. Lahens, N. F. et al. Time-specific associations of wearable sensor-based cardiovascular and behavioral readouts with disease phenotypes in the outpatient setting of the Chronic Renal Insufficiency Cohort. Digit Health 8, 20552076221107903 (2022).

42. Zhang, S. L. et al. A circadian clock regulates efflux by the blood-brain barrier in mice and human cells. Nat Commun 12, 617 (2021).

43. Zielinski, T., Moore, A. M., Troup, E., Halliday, K. J. & Millar, A. J. Strengths and Limitations of Period Estimation Methods for Circadian Data. PLOS ONE 9, e96462 (2014).

44. Dobin, A. et al. STAR: ultrafast universal RNA-seq aligner. Bioinformatics 29, 15–21 (2013).

45. Yates, A. D. et al. Ensembl 2020. Nucleic Acids Research 48, D682–D688 (2020).

46. Weger, B. D. et al. Systematic analysis of differential rhythmic liver gene expression mediated by the circadian clock and feeding rhythms. Proceedings of the National Academy of Sciences 118, e2015803118 (2021).

47. Brooks, T. G. et al. Nitecap: An Exploratory Circadian Analysis Web Application. J Biol Rhythms 37, 43–52 (2022).

48. Qin, Q. et al. Lisa: inferring transcriptional regulators through integrative modeling of public chromatin accessibility and ChIP-seq data. Genome Biol 21, 32 (2020).

